# Landscapes and bacterial signatures of mucosa-associated intestinal microbiota in Chilean and Spanish patients with inflammatory bowel disease

**DOI:** 10.1101/2021.01.04.425212

**Authors:** Nayaret Chamorro, David A. Montero, Pablo Gallardo, Mauricio Farfán, Mauricio Contreras, Marjorie De la Fuente, Karen Dubois, Marcela A. Hermoso, Rodrigo Quera, Marjorie Pizarro-Guajardo, Daniel Paredes-Sabja, Daniel Ginard, Ramon Rosselló-Móra, Roberto Vidal

## Abstract

Inflammatory bowel diseases (IBD), which include ulcerative colitis (UC) and Crohn’s disease (CD), cause chronic inflammation of the gut, affecting millions of people worldwide. IBD have been frequently associated with an alteration of the gut microbiota, termed dysbiosis, which is generally characterized by an increase in abundance of Proteobacteria such as *Escherichia coli,* and a decrease in abundance of Firmicutes such as *Faecalibacterium prausnitzii* (an indicator of a healthy colonic microbiota). The mechanisms behind the development of the IBD and the dysbiosis are incompletely understood. Using samples from colonic biopsies, we studied the mucosa-associated intestinal microbiota in Chilean and Spanish patients with IBD. In agreement with previous studies, microbiome comparison between IBD patients and non-IBD controls indicated that dysbiosis in these patients is characterized by an increase of pro-inflammatory bacteria (mostly Proteobacteria) and a decrease of commensal beneficial bacteria (mostly Firmicutes). Notably, bacteria typically residing on the mucosa of healthy individuals were mostly obligate anaerobes, whereas in the inflamed mucosa an increase of facultative anaerobe and aerobic bacteria was observed. We also identify potential co-occurring and mutually exclusive interactions between bacteria associated with the healthy and inflamed mucosa, which appear to be determined by the oxygen availability and the type of respiration. Finally, we identify a panel of bacterial biomarkers that allow the discrimination between eubiosis from dysbiosis with a high diagnostic performance (96% accurately), which could be used for the development of non-invasive diagnostic methods. Thus, this study is a step forward toward understanding the landscapes and alterations of mucosa-associated intestinal microbiota in patients with IBD.

## INTRODUCTION

The intestinal microbiota plays a key role in human health, providing important metabolic functions, stimulating the immune system, acting as a barrier to pathogenic organisms, and regulating body composition [1, 2]. Changes in microbiota structure or dysbiosis have been associated with alterations in diet [3], chronic stress [4], antibiotic use and a number of gastrointestinal disorders [5–10], such as the inflammatory bowel disease (IBD) [11, 12]. Within IBD phenotypes, ulcerative colitis (UC) and Crohn’s disease (CD) have a clinical impact worldwide, with UC characterized by inflammation of the rectum, extending diffusely towards the colon, whereas CD is characterized by systemic inflammation and ulcers affecting any part of the gastrointestinal tract and thickening of the intestinal wall. Since 1990, the incidence of IBD has increase in Africa, Asia and South America [13]; although no data are available in Chile, clinical experience shows an increase in recent years in the number of consultations by IBD patients [14, 15].

While the etiological causes of UC and CD remain unidentified, several factors are known to increase susceptibility, including: i) genetic polymorphisms [16–18]; ii) altered immune response [19, 20] iii) environmental factors [21–23]; and (iv) alteration of the gut microbiota composition [24–29].

In general, studies of the gut microbial composition of patients with UC and CD have shown an increase in the phylum Proteobacteria and a decrease in the phylum Firmicutes [30, 31]. Most of these studies have mainly analyzed stool samples. However, many differences between the microbial communities from fecal and mucosal samples have been reported, indicating that fecal microbial communities do not accurately represent the local communities that live in specific regions of the gut (colon, ileus and small intestine) [20, 32–34]. In addition, most of these studies are based on the massive sequencing of amplicons of the 16S rRNA gene and the clustering of reads into operational taxonomic units (OTUs), using short sequences of variable regions of the gene [35, 36]. The short size of the amplicons (<300 nt) only allows the identification up to the family level or in some cases genus.

In the present study, we investigated the mucosa-associated intestinal microbiota in Chilean and Spanish patients with IBD. For this, we amplified the 16S rRNA gene and obtained sequence libraries from colonic mucosa biopsies using high-quality reads of more than 300 nucleotides for the bacterial affiliation process. In addition, we used the OPU (Operational Phylogenetic Unit) approach for taxonomic assignment, as this allows for better bacterial identification, in most cases reaching the species level [37]. OPU analysis is not based on strict identity thresholds; instead, sequences are affiliated with a phylogenetic tree using the parsimony algorithm followed by manual supervision of the tree to design meaningful phylogenetic units [38]. Since identification is based on phylogenetic inference, OPUs are based on the genealogical signal of the sequences, which minimizes the influence of errors and size differences. An OPU is the smallest monophyletic group of sequences containing OTU representatives together with the closest reference sequence, including the sequence of a type strain when possible [39]. To date, there have been no reports on the microbiome of patients with UC at the OPU level.

Our results indicate that dysbiosis is not a common feature among IBD patients. However, in IBD patients where gut dysbiosis was present, we found an increase in pro-inflammatory bacteria (mostly Proteobacteria) and a decrease in beneficial commensal bacteria (mostly Firmicutes). We also identified potential co-occurring and mutually exclusive interactions between bacteria associated with the healthy and inflamed mucosa, which appear to be determined by oxygen availability and the type of respiration. Importantly, these results were consistent with the “Oxygen Hypothesis”, which states that chronic inflammation induces increased oxygen levels in the gut, leading to an imbalance between obligate and facultative anaerobes [28]. Finally, we identified bacterial biomarkers that could be used for the development of non-invasive diagnostic methods, such as real-time PCR. A future goal is that an early detection of changes in the gut microbiota could allow the initiation of IBD treatment or preventive measures.

## RESULTS

Here we focused on two independent cohorts of patients with IBD from Chile and Spain. Both cohorts were adults and included patients diagnosed with UC or CD. In addition, control individuals (CTL; Non-IBD controls) who underwent colonoscopy due to a family history of colon cancer were included. All samples were colonic mucosal biopsies. Patients who received antibiotic treatment within one month prior to the colonoscopy were excluded. The Chilean cohort included 20 and 21 patients with UC and CD, respectively, and 5 control individuals. The Spanish cohort was previously reported by Vidal et al., 2015 [37], of which we included 13 patients with CD and also, 7 control individuals. The clinical features of the patients are described in Table S1. For both cohorts, microbial composition in colonic biopsies were assessed by DNA extraction followed by 16S rRNA gene pyrosequencing, as described in the Materials and Methods section.

Pyrosequencing of Chilean samples generated 331,677 reads, which together with the Spanish samples generated a total of 483,818 reads with a mean of 5,506 reads per sample (Table S2). The average read length for 16S rRNA sequences was 649 bp. The OTUs from Chilean samples were clustered and inserted into the tree by Vidal et al., 2015 [37]. OTUs that were not clustered as part of known OPUs were re-evaluated and designated anew. This resulted in a total of 608 OPUs, with a mean of 88 OPUs per sample (Table S2). Notably, our results show that the OPU rarefaction curves approached saturation with a significantly lower number of reads than necessary for the OTU curves, indicating an overestimation of taxonomic units (diversity) when using a traditional OTU approach (Figure 1A).

**FIGURE 1.**
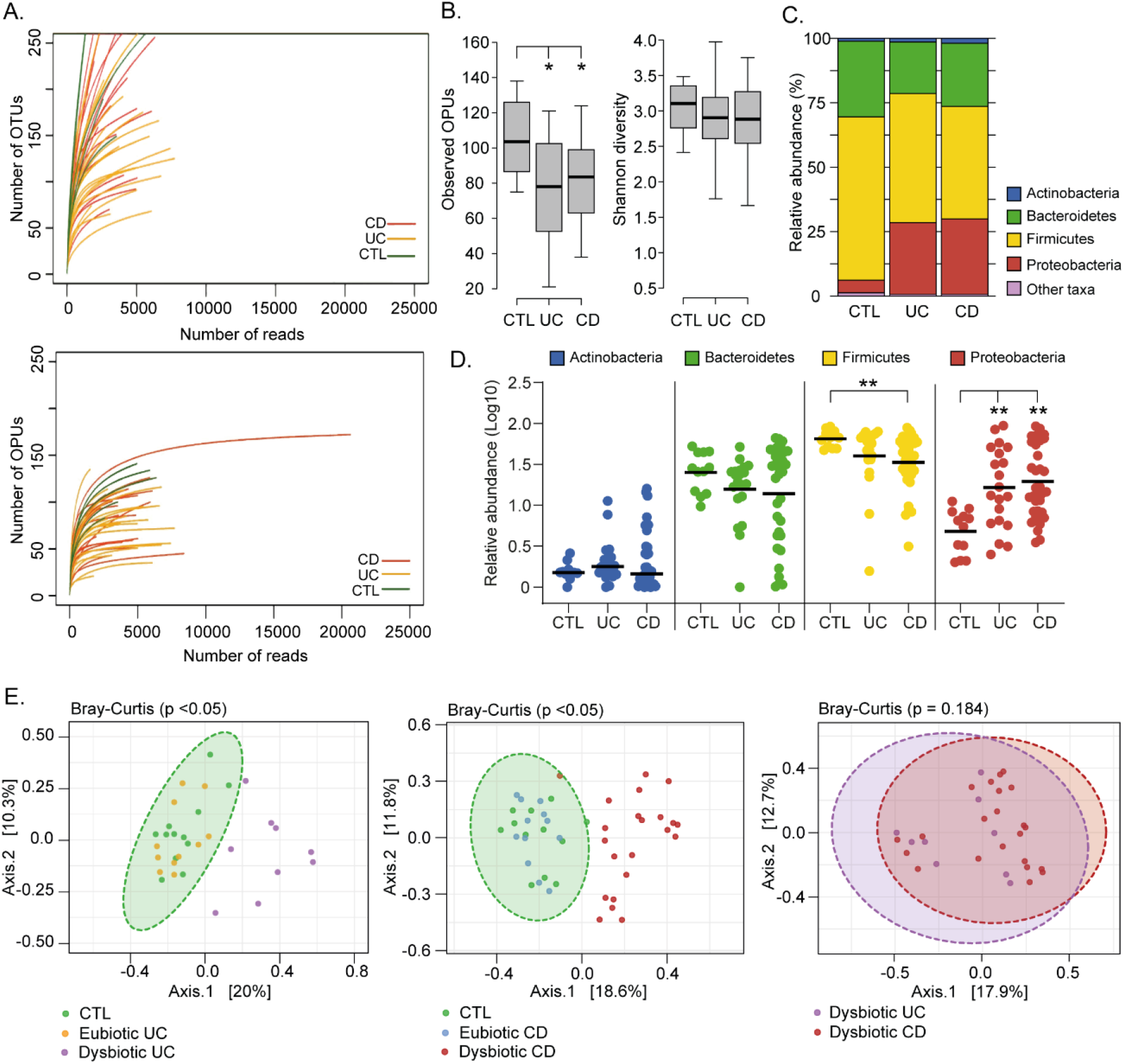
Landscape of mucosa-associated intestinal microbiota of Chilean and Spanish patients with IBD. **(A)** Rarefaction curves based on OTUs and OPUs detected in colonic biopsies from patients with ulcerative colitis (UC; orange lines) and Crohn’s Disease (CD; red lines), and control individuals (CTL; green lines). **(B)** Alpha diversity between patients and control individuals determined by the observed OPUs (Richness) and the Shannon index (Diversity). Significance: * *p* < 0.05, Kruskal-Wallis & Dunn’s tests. **(C)** Relative abundances (%) of the most prevalent phyla. Phyla with abundances < 1% are represented as other taxa. **(D)** Comparison of the relative abundances (Log10) of the four most common phyla. Each data point corresponds to a sample and horizontal lines to the means. Significance: ** *p* < 0.005, Kruskal-Wallis & Dunn’s tests. **(E)** A set of principal coordinate analysis (PCoA) plots based on Bray-Curtis distances showing the overall composition (beta diversity) of the microbiota in patients and controls. For a better visualization, individual PCoA plots are shown for the analysis of UC patients versus controls (left panel), CD patients versus controls (middle panel) and dysbiotic UC versus dysbiotic CD patients. Each data point corresponds to a sample, which is colored according to the disease phenotype and the microbiota status (dysbiotic or eubiotic). Ellipses represent a 95% CI around the cluster centroid.

### Gut microbial dysbiosis is not a common feature among IBD patients

Differences in the gut microbiota of IBD patients compared to healthy individuals have been widely reported [40–43]. Among IBD patients there may be several degrees and profiles of dysbiosis, which have been correlated with the phenotype and severity of the disease [35, 37, 44], and gastrointestinal surgery [45, 46]. Therefore, we first assessed the landscape, richness, and diversity of the mucosa-associated intestinal microbiota among patients and non-IBD controls. In general, we found that UC and CD patients have significantly lower richness (Observed OPUs) than the controls. However, no significant differences in the alpha diversity (Shannon diversity) were observed (Fig. 1B).

Four major bacterial phyla (Firmicutes, Bacteroidetes, Proteobacteria and Actinobacteria) dominate the gut microbiota of humans [33]. Accordingly, almost all OPUs identified were affiliated with one of these phyla. In general, we observed that UC and CD patients have a lower abundance of Firmicutes and a higher abundance of Proteobacteria (Fig. 1C and 1D). Interestingly, a principal coordinate analysis (PCoA) based on the Bray-Curtis dissimilarity showed that only some patients have a significant shift in beta diversity (dysbiosis) compared to the controls (Fig. 1E). This result suggests that some patients (dysbiotic UC and CD patients) but not all (eubiotic UC and CD patients) have an alteration in their mucosa-associated intestinal microbiota. Of note, there were no significant differences in beta diversity between dysbiotic patients that clustered away from control individuals, suggesting they have a similar imbalance in their microbiota.

Based on the PCoA analysis and the IBD phenotype, patients were subdivided into four groups: UC1, UC2, CD1 and CD2 (Table S1). Consistent with this subclassification, hierarchical clustering using the relative abundances of the 608 OPUs across all samples showed that UC1, CD1 and controls clustered together but away from UC2 and CD2 (Fig. 2A). Moreover, relative abundances per patient at the phylum level clearly showed that while UC1, CD1 and controls have a similar taxonomic composition, UC2 and CD2 have a dysbiosis characterized by an increase in Proteobacteria and a decrease in Firmicutes (Fig. 2B). As expected, UC1 and CD1 showed a richness and diversity similar to the controls. By contrast, UC2 and CD2 had significantly lower richness than the controls, but only CD2 showed a lower alpha diversity (Fig. 2C). Thus, collectively these results indicate variability in the mucosa-associated intestinal microbiota among IBD patients: while some patients have a dysbiotic microbiota, others have a microbiota composition similar to that of the non-IBD controls.

**FIGURE 2.**
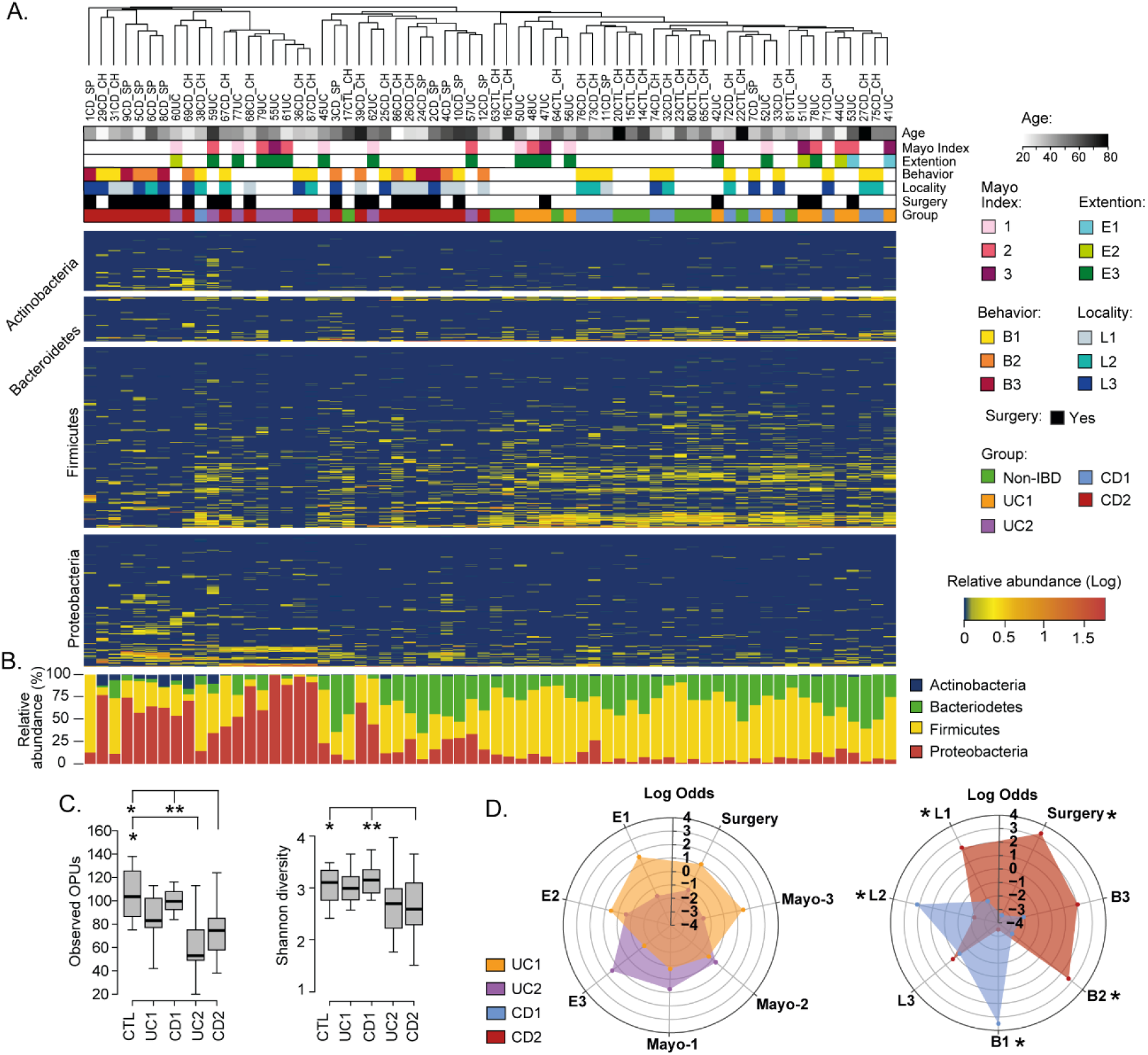
Compositional differences in the mucosa-associated intestinal microbiota among IBD patients. **(A)** Clustering of patients and controls and heatmap of OPU abundances. Patients (Columns) were grouped based on a hierarchical cluster analysis using relative abundances of 608 OPUs. OPUs of the four most abundant phyla are shown in the rows, and their relative abundances are shown on a log scale according to the legend. The clinical and demographic features of the patients are shown at the top and are colored according to the legend. Note that in general, patients belonging to the UC2 and CD2 groups clustered together and away from patients in the UC1 and CD1 groups and the controls (Non-IBD). **(B)** Relative abundances (%) per patient of the four most abundant phyla. **(C)** Alpha diversity between IBD groups (UC1, UC2. CD1, and CD2) and control individuals determined by the observed OPUs (Richness) and the Shannon index (Diversity). Significance: * *p* < 0.05, ** *p* < 0.005, Kruskal-Wallis & Dunn’s tests. **(D)** Radar chart showing association of IBD groups with clinical features. Pairwise association between patient groups and clinical features was performed in contingency tables by odds ratios. Significance: * *p* < 0.05, Pearson’s chi-squared test or Fisher’s exact test. The figure was prepared using the Plotly package [47] in R [48].

### Dysbiosis in Crohn’s patients is correlated with disease severity

We then asked whether patient groups correlated with demographic and clinical variables. Regarding demographic features, no significant correlations were found with respect to age, sex, or origin of the patients (not shown). Similarly, there were no significant correlations of the UC1 and UC2 groups with clinical variables. By contrast, while the CD1 group correlated with colonic location (L2) and an inflammatory disease behavior (B1), the CD2 group correlated with ileal location (L1), the stricturing (B2) and penetrating (B3) phenotypes, and surgery (Fig. 2D). Thus, patients in the CD2 group were associated with an advanced stage of the disease, which suggests a clinically relevant link between gut microbiota dysbiosis and disease severity.

### Dysbiosis in IBD is characterized by an increase in pathobionts and a decrease in anti-inflammatory commensal bacteria

Although it is not known whether dysbiosis is a cause or a consequence of IBD, the identification of specific taxa and bacteria (biomarkers) associated with these patients may contribute to understanding the dynamics of this disease (progression or remission), as well as the development of diagnostic aids [49]. Therefore, we sought to characterize the variation in the gut microbiota at different taxonomic levels among the IBD patients and controls. We found that the relative abundances at the phylum and class levels were similar between the UC1, CD1 and control groups, further confirming that these IBD patients have a non-dysbiotic gut microbiota (eubiotic status) (Fig. 3). Conversely, in the UC2 and CD2 groups, the phylum Firmicutes was decreased, mainly attributed to a lower abundance in Clostridia, while the phylum Proteobacteria was increased, mainly attributed to a higher abundance in Alpha- and Gammaproteobacteria (Fig. 3A and 3B).

**FIGURE 3.**
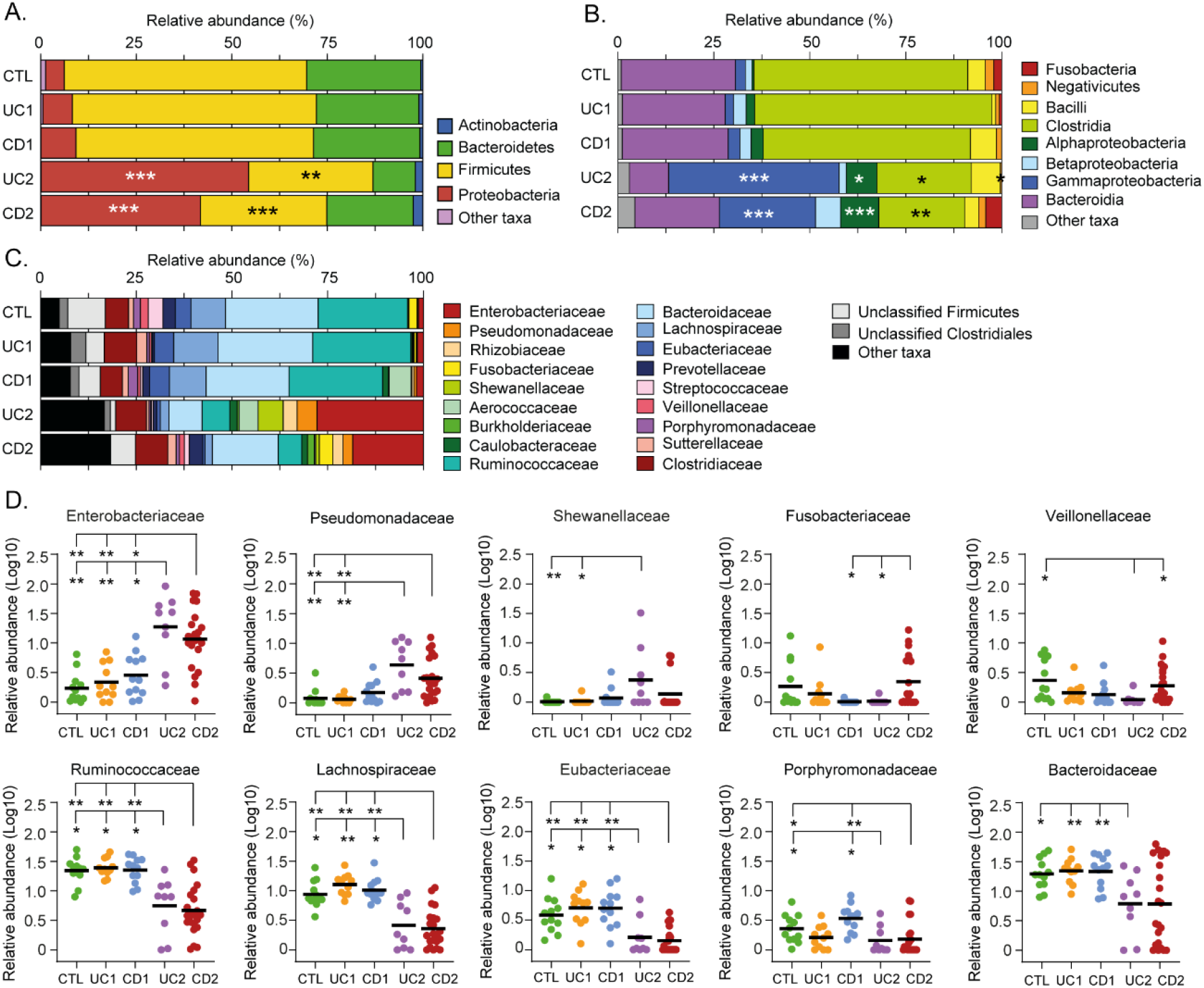
Phylogenetic composition of bacterial taxa at the phylum, class, and family level among IBD patients and non-IBD controls. Relative abundances (%) of common bacterial taxa (> 1% abundance) at phylum **(A)**, class **(B)**, and family **(C)** levels. **(D)** For a better analysis and visualization, comparisons in the relative abundances at the family level between groups were performed using Log-transformed values. Each data point corresponds to a sample and horizontal lines to the means. Only taxa in which significant differences were found are shown. Significance: * *p* < 0.05, ** *p* < 0.005, *** *p* < 0.0005, Kruskal-Wallis & Dunn’s tests.

At the family level, we found that the UC2 and CD2 groups had a greater abundance of Enterobacteriaceae and Pseudomonadaceae. Additionally, Shewanellaceae was increased in UC2. The shift in the gut microbiota composition in the UC2 and CD2 groups was also characterized by a decreased abundance of Ruminococcaceae, Lachnosporiraceae, Eubacteriaceae, Porphyromonadaceae and Bacteroidaceae (Fig. 3C and 3D). On the other hand, differences between the UC2 and CD2 groups were only observed in Fusobacteriaceae and Veillonellaceae, with both taxa being decreased in the UC2 group. Hence, the dysbiotic profile of UC2 and CD2 patients was in general very similar.

To further investigate the microbial imbalance in the IBD, we used the LEfSe algorithm to identify which OPUs were differentially abundant in the dysbiotic patients (UC2 and CD2 groups) compared to the eubiotic patients (UC1 and CD1 groups) and controls. As a result, 24 OPUs were found to be significantly increased in dysbiotic patients, including several potential pathogens (pathobionts) such as *Escherichia coli, Klebsiella oxytoca, Ruminococcus gnavus, Enterococcus faecalis,* several *Rhizobium, Pseudomonas and Clostridium* species, and members of the tribe *Proteeae (Providencia/Morganella)* (Fig. 4A, Table S3). By contrast, 25 OPUs were found to be decreased, including several symbionts known to provide important functions for gut health, mainly through the production of butyrate [50, 51]. For instance, butyrate producers such as *Faecalibacterium prausnitzii*, *Blautia*, *Gemmiger formicilis*, *Eubacterium rectale, Ruminococcus torques, Roseburia inulinivorans and Coprococcus catus* were significantly decreased. Other commensal bacteria such as *Alistipes* and *Dorea* were also decreased in these patients. Thus, collectively these results indicate that the dysbiosis in these IBD patients is characterized by an increase in pathobionts and a decrease in beneficial commensal bacteria.

**FIGURE 4.**
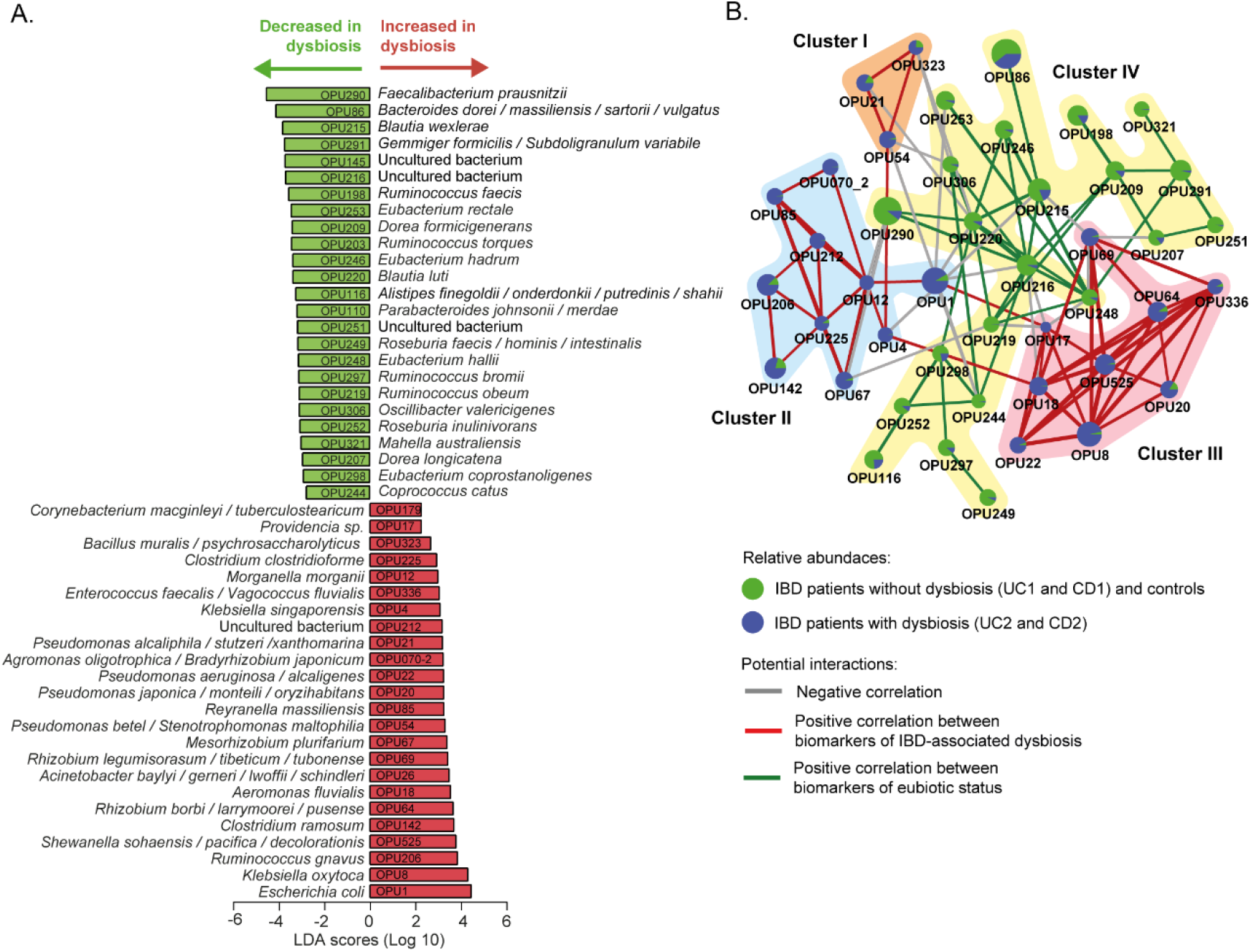
Differently abundant OPUs in dysbiotic IBD patients and their coexistence networks. **(A)** The LEfSe algorithm with a linear discriminant analysis (LDA) score >2 enabled the identification of OPUs that differed significantly in abundance between the dysbiotic patients (UC2 and CD2 groups) and eubiotic patients (UC1 and CD1) and non-IBD controls **(B)** Correlation networks of differently abundant OPUs. This analysis was performed using SparCC [52] (see methods). The nodes (pie charts) represent OPUs, with size reflecting the relative abundance in IBD patients and controls, as described in the legend. The links between nodes correspond to significant interactions (positive or negative correlations), with line width reflecting the strength of the correlation. Clusters (I to IV) of highly correlated OPUs are indicated. Unconnected nodes were omitted.

### Pathobionts increased in dysbiotic IBD patients have co-occurring relationships and do not co-exist with the core bacteria of the eubiotic state

In the context of IBD, unraveling potential interactions between bacteria associated with a healthy or an affected mucosa can contribute to understanding how the microbiota responds and adapts to an inflammatory environment. The correlation network analysis has been used to predictively model the interplay between the microbiota and the environment [53]. Consequently, in the following we explore the potential interactions among the OPUs that were differentially abundant in IBD patients using the SparCC algorithm [52].

Notably, this analysis revealed that pathobionts increased in dysbiosis form three clusters (I, II and III) of positive co-occurring relationships (Fig 4B). In particular, Cluster I (OPU21, OPU54 and OPU323) and Cluster III (OPU8, OPU17, OPU18, OPU20, OPU22, OPU64, OPU69, OPU336 and OPU525) include facultative anaerobic and aerobic bacteria, and Cluster II (OPU1, OPU4, OPU12, OPU67, OPU70-2, OPU85, OPU142, OPU206, OPU212 and OPU225) includes obligate anaerobic, facultative anaerobic and aerobic bacteria (Table S3). Furthermore, these three clusters have negative correlations (mutual exclusivity) with Cluster IV, which is formed by obligate anaerobic bacteria that were significantly abundant in patients without dysbiosis and the controls. Overall, this result indicates that bacteria of Clusters I and III and the majority of Cluster II can survive in a niche with a high level of oxygen. Therefore, oxygen levels at the intestinal mucosa could be an environmental characteristic that determine the type of relationship (co-existence or mutual exclusivity) between these bacteria.

### Dysbiosis in IBD patients is characterized by a shift from obligate to facultative anaerobes and to aerobic bacteria

The healthy intestinal mucosa has low levels of oxygen and thereby allows the survival and establishment of bacterial communities of obligate anaerobes. By contrast, dysbiosis in IBD is characterized by an increase in facultative anaerobes from the phylum Proteobacteria. Furthermore, it has been hypothesized (Oxygen Hypothesis) that chronic inflammation leads to an increased oxygen levels in the gut, which in turn creates a microenvironment that favors facultative anaerobes or even aerobic bacteria [28]. In light of these observations, we evaluated the oxygen hypothesis by investigating the microbial respiration of mucosa-associated intestinal microbiota among the IBD patients. Interestingly, consistent with the oxygen hypothesis and previous studies [54, 55], we found that the IBD patients with dysbiosis (UC2 and CD2) have a decreased abundance of obligate anaerobes and an increased abundance of facultative anaerobes and aerobic bacteria. Thus, this result, together with the correlation network analysis, strongly suggests a key role for oxygen in the intestinal dysbiosis of IBD patients.

### Bacterial biomarkers make it possible to discriminate dysbiosis and eubiosis in IBD patients and between UC and CD patients with dysbiosis

Currently, IBD has no clear etiology, and due to non-specific symptoms, diagnosis in some patients can be delayed or missed [56]. In this sense, diagnostics is a promising application of the microbiota association studies. It is important to note that instead of classifying IBD patients based on the overall gut microbiota composition, the use of a small set of bacterial biomarkers would be more feasible and economical in clinical practice. Moreover, since dysbiosis is not a common feature among IBD patients, we rationalize that as a first approach, discriminating dysbiosis from eubiosis in these patients could be a valuable diagnostic aid.

Therefore, we evaluated the 49 differentially abundant OPUs as biomarkers using the ROC curve analysis. This analysis showed that some of these OPUs could discriminate dysbiosis from eubiosis but with a fair to poor diagnostic performance (Figure S1). The best performance as biomarkers were achieved by *Faecalibacterium prausnitzii* (OPU290), *Blautia luti* (OPU220), *Coprococcus catus* (OPU244), *Eubacterium hadrum* (OPU246) and *Escherichia coli* (OPU1), with an area under the curve (AUC) ranging from 0.87 to 0.94.

Next, to improve the diagnostic performance achieved by individual biomarkers, we used a combinatorial panel of OPUs and implemented five machine learning classification methods (Neural Network, Naïve Bayes, Logistic regression, Random Forest, and Support Vector Machines) (see methods). For this, we selected ten biomarkers of eubiosis and ten biomarkers of dysbiosis based on their AUC. As a result, we found that this panel of twenty biomarkers enabled dysbiosis to be distinguished from eubiosis with a high discriminatory power, with an AUC ranging from 0.96 to 0.99 (Fig. 6, Table 1). In particular, the Neural Network and Naïve Bayes models were the best performing classifiers, both with an overall accuracy of 96% (Table 1).

**TABLE 1.** Cross-validation of bacterial biomarkers and machine learning methods for diagnostic aids in IBD

| Method | Average<br>AUC | CA | IBD-Dysbiotic profile |  |  | Eubiotic profile |  |  |
| --- | --- | --- | --- | --- | --- | --- | --- | --- |
|  |  |  | F1 | Precision | Recall | F1 | Precision | Recall |
| TOP 20 indicator OPUs (IBD Dysbiosis vs. Eubiosis) |  |  |  |  |  |  |  |  |
| SVM | 0.97 | 0.92 | 0.92 | 0.96 | 0.87 | 0.93 | 0.89 | 0.97 |
| Random Forest | 0.96 | 0.88 | 0.87 | 0.89 | 0.84 | 0.89 | 0.87 | 0.91 |
| Neural Network | 0.99 | 0.96 | 0.95 | 1.00 | 0.903 | 0.96 | 0.92 | 1.00 |
| Naïve Bayes | 0.99 | 0.96 | 0.95 | 0.97 | 0.94 | 0.96 | 0.94 | 0.97 |
| Logistic Regression | 0.97 | 0.89 | 0.89 | 0.90 | 0.87 | 0.90 | 0.89 | 0.91 |
| Method | Average<br>AUC | CA | UC-Dysbiotic profile |  |  | CD-Dysbiotic profile |  |  |
|  |  |  | F1 | Precision | Recall | F1 | Precision | Recall |
| Top 11 indicator OPUs (UC2 – CD2 groups) |  |  |  |  |  |  |  |  |
| SVM | 0.68 | 0.74 | 0.429 | 0.60 | 0.33 | 0.83 | 0.77 | 0.91 |
| Random Forest | 0.82 | 0.68 | 0.38 | 0.43 | 0.33 | 0.78 | 0.75 | 0.82 |
| Neural Network | 0.75 | 0.74 | 0.56 | 0.56 | 0.56 | 0.82 | 0.82 | 0.82 |
| Naïve Bayes | 0.83 | 0.77 | 0.72 | 0.56 | 1.00 | 0.81 | 1.00 | 0.68 |
| Logistic Regression | 0.80 | 0.71 | 0.53 | 0.50 | 0.56 | 0.79 | 0.81 | 0.77 |

SVM, Support Vector Machine. AUC, Area under the curve. CA, Classification accuracy measures the ratio of the correct predictions to the total number of instances evaluated. F1-measure denotes the harmonic mean between recall and precision values, where an F1 score reaches its best value at 1 and the worst score at 0.

Finally, because the microbiota composition between the UC and CD patients with dysbiosis was remarkably similar, we sought to clarify whether bacterial biomarkers can discriminate between these groups of patients. For this, we implemented the same prediction pipeline shown above, that is, the LEfSe algorithm followed by a ROC curve analysis and machine learning classification methods. In this case, we selected a panel of four biomarkers of UC patients with dysbiosis and seven biomarkers for CD patients with dysbiosis (Fig. 6C). By using these eleven biomarkers, the Naïve Bayes model was the best classifier, achieving an AUC and accuracy of 0.83 and 77%, respectively (Fig. 6D, Table 1). Collectively, these results highlight the potential use of these bacterial markers as a diagnostic aid in IBD.

## DISCUSSION

Several studies have investigated the gut microbiota composition in patients with IBD, but most have mainly analyzed stool samples [42–44]. Fecal samples are readily available and easily collected, allowing easy longitudinal sampling within individuals with no major methodological complexities. However, fecal microbial communities do not accurately represent the mucosa-associated microbiota that live in the gastrointestinal tract (e.g. colon, ileus and small intestine) [32–34]. Moreover, characterization of the microbiota at mucosal lesion sites provides relevant insights and more accurate conclusions regarding taxonomic composition and dysbiosis in gastrointestinal disorders such as IBD [23, 57]. For instance, Gevers et al. [36] investigated the composition of the microbiota in IBD patients using different sample types (stool samples and tissue biopsies of the ileum and rectum) and found that the rectal mucosa-associated microbiota has the potential for an early and accurate diagnosis of CD. By contrast, the composition of the fecal microbiota was less informative with a poor diagnostic performance in IBD.

In this work, we determined the landscapes and bacterial signatures of mucosa-associated microbiota in two independent cohorts of Chilean and Spanish patients diagnosed with IBD. For this, we used the Operational Phylogenetic Unit (OPU) concept for taxonomic assignment [37], which allowed us to reach the species level with a lower number of reads than needed for the traditional OTU-based approach (Fig. 1A). Our results showed that while some IBD patients had an imbalance in the mucosa-associated microbiota, other patients had a microbial composition that was similar to that of non-IBD controls (Fig. 1B-E). Hence, dysbiosis was not a common feature among the IBD patients, and thus, we decided to subdivide them (based on the microbiota composition and the IBD phenotype) into four groups: two eubiotic groups (UC1 and CD1) and two dysbiotic groups (UC2 and CD2). Importantly, this subclassification was consistent with the hierarchical clustering of patients in which the UC1, CD1 and non-IBD controls clustered together but away from UC2 and CD2 (Fig. 2A). Our results showed that the dysbiosis in UC2 and CD2 was characterized by an increase in Proteobacteria and a decrease in Firmicutes (Fig. 2B and Fig. 3). Previous studies have also shown the same alteration in the abundance of these phyla in the mucosa-associated microbiota of IBD patients [30, 58, 59]. Moreover, differences between the UC2 and CD2 groups were only observed at the family level, with a decreased abundance of Fusobacteriaceae and Veillonellaceae in the UC2 group (Fig. 3D). Despite the similarities in the microbial composition of UC2 and CD2 groups, dysbiosis was correlated with disease severity only in the latter group (Fig. 2D).

Several bacteria (pathobionts) that were found to be increased in the mucosal samples of dysbiotic IBD patients are known to have the potential to exacerbate inflammation (Fig. 4A and Table S3). For instance, pathogenic *E. coli* strains with the ability to adhere and invade the intestinal mucosa have been isolated with a high frequency from biopsies of CD patients [26, 27]. *Klebsiella oxytoca* causes antibiotic-associated hemorrhagic colitis [60, 61]. *Ruminococcus gnavus* produces an inflammatory polysaccharide and an increased abundance of this bacterium in IBD patients has been linked to an increase in disease activity [62–64]. *Enterococcus faecalis* metalloprotease GelE disrupts the epithelial barrier and increased intestinal inflammation in interleukin-10 knockout mice [65–67]. An increased abundance of *Rhizobium* spp has been shown in CD patients with recurrence after surgery compared to those who remain in remission [46]. Similarly, an increased abundance of *Pseudomonas* spp has been reported in IBD patients [44, 68]. Moreover, *Pseudomonas* spp. can attach to intestinal epithelial cells and deliver toxins to host cells through a type 3 secretion system, thereby causing epithelial cell damage [69, 70]. *Clostridium ramosum* has been reported to be increased in pediatric patients with CD and induces ulcerative colitis in the DSS-colitis mouse model [71, 72].

The dysbiosis in the IBD patients was also characterized by a decreased abundance of several commensal bacteria with anti-inflammatory properties, including several butyrate producers such as *Faecalibacterium prausnitzii, Blautia, Gemmiger formicilis, Eubacterium rectale, Ruminococcus torques, Roseburia inulinivorans and Coprococcus catus* [73]. This result is in agreement with previous studies showing a reduced abundance of multiple butyrate-producing bacteria in dysbiotic IBD patients [41, 45, 74]. Butyrate is a short-chain fatty acid that plays an important role in maintaining the integrity of colonic mucosa and regulating cell proliferation and differentiation [50, 51]. Studies on biopsies from patients with CD cultured with butyrate showed a dose-dependent decrease in the expression of proinflammatory cytokines [75]. In particular, *Faecalibacterium prausnitzii* is known to be a core bacterium of the gut microbiota of healthy adults, representing around 5% of the total bacterial population. This bacterium plays an important role in butyrate production and has a strong anti-inflammatory effect [73]. Many studies have shown a decreased abundance of this bacterium in IBD patients, this reduction being a bacterial signature of dysbiosis as well as the severity and activity of the disease [24, 76, 77]. Additional studies are needed to investigate whether the increased abundance of the pathobiont and the depletion of butyrate-producing bacteria may trigger pro-inflammatory signals that contribute to the development or progression of the IBD.

An interesting finding of this study was the identification of potential co-occurring relationships between pathobionts associated with the inflamed mucosa (Fig. 4B). In turn, these pathobionts showed mutually exclusive relationships with the core bacteria of the healthy mucosa. In an ecological context, a cluster network of highly correlated organisms could be supported by synergistic, related, or shared biological function, including metabolism, respiration, trophic chains, specific bacterial features (immune evasion, aggregation / biofilm formation) and habitat characteristics (pH, salinity, immune recognition, mucins, among others). In line with this idea, we found that that most of the OPUs that form Clusters I, II and III are facultative anaerobic or aerobic bacteria. Conversely, the OPUs of Cluster IV are obligate anaerobic bacteria (Table S3). Thus, pathobionts belonging to Clusters I, II and III could be favored by the inflammation and thickness reduction of the mucus layer, which leads to a higher oxygen level in the intestinal epithelium. Moreover, this result is consistent with the study by Hughes et al., 2017 [54], in which formate oxidation and oxygen respiration were identified as metabolic signatures for inflammation-associated dysbiosis.

Importantly, we observed an increased abundance of facultative anaerobes and aerobic bacteria in dysbiotic IBD patients (Fig. 5), which is consistent with the oxygen hypothesis. This hypothesis states that chronic inflammation in IBD patients leads to a greater release of hemoglobin (which carries oxygen) and reactive oxygen species to the intestinal lumen. Thus, the increase in oxygen levels causes a disruption in anaerobiosis that confers selective advantages to facultative anaerobes and aerobes, allowing them to become more competitive and able to overgrow [28].

**FIGURE 5.**
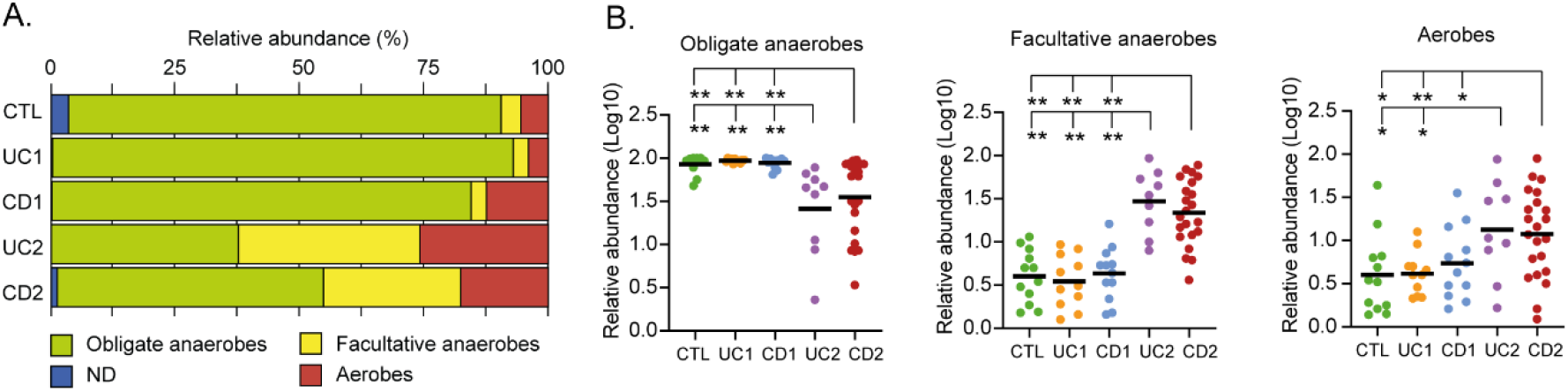
Microbial respiration of mucosa-associated intestinal microbiota among the IBD patients. **(A)** Relative abundances (%) of OPUs classified as obligate anaerobes, facultative anaerobes, or aerobes. **(B)** Differences in the relative abundances (Log10) of obligate anaerobes, facultative anaerobes, or aerobes. Each data point corresponds to a sample and horizontal lines to the means. Significance: * *p* < 0.05, ** *p* < 0.005, Kruskal-Wallis & Dunn’s tests. ND: Unclassified bacteria with unknown respiration type.

Cost-effective, rapid, and reproducible biomarkers would be helpful for patients and clinicians in the diagnosis of IBD. Several bacterial biomarkers for IBD diagnosis have been evaluated in previous studies with promising results [42, 56, 78–80]. The panel of bacterial biomarkers identified in the current study made it possible to discriminate dysbiosis from eubiosis in IBD with a high discriminatory power (96% accurately) (Fig. 6A). Likewise, our bacterial biomarkers discriminated between the dysbiotic UC patient from the dysbiotic CD patients, although with a lower diagnostic performance (77% accurately) (Fig. 6B). The cross-validation of these bacterial biomarkers in different cohorts will confirm their potential use as a diagnostic aid. Finally, these biomarkers could be combined with imaging techniques and calprotectin levels to improve the diagnosis of IBD, thereby facilitating clinical decision-making.

**FIGURE 6.**
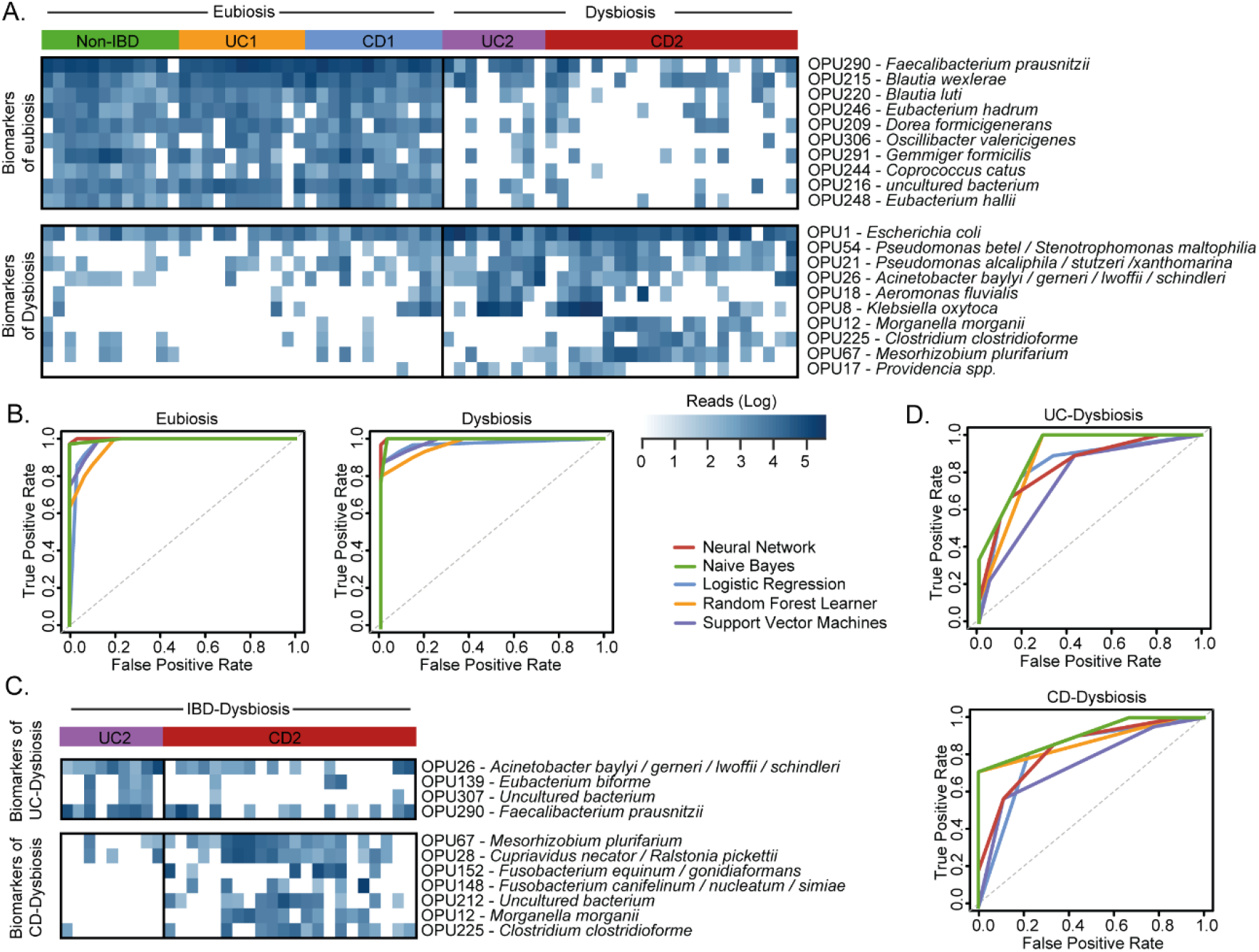
Evaluation of OPUs as biomarkers for IBD. **(A)** Heatmap showing the number of reads (log scale) of the best 20 indicator OPUs for dysbiosis and eubiosis in IBD. Rows represent OPUs and columns the patients ordered by groups. **(B)** Evaluation of five machine learning models and the above panel of twenty indicator OPUs to discriminate dysbiosis from eubiosis in the IBD patients. **(C)** Heatmap showing the number of reads (log scale) of the best 11 indicator OPUs for the UC (UC2) and CD (CD2) patients with dysbiosis. Rows represent OPUs and columns the patients ordered by groups. **(D)** Evaluation of five machine learning models and the above panel of 11 indicator OPUs to discriminate UC2 patients from CD2 patients. The diagnostic performance of each classifier model is represented by ROC curves and the area under the curve is indicated in Table 1.

## MATERIALS AND METHODS

### Patients

Two independent cohorts of patients with IBD from Chile and Spain were included in this study (Table S1). Both cohorts were adults and included 20 Chilean patients diagnosed with ulcerative colitis (UC_CH), 21 Chilean patients diagnosed with Crohn’s disease (CD_CH) and 5 Chilean non-IBD control individuals (CTL_CH) who underwent a colonoscopy due to a family history of colon cancer. The Spanish cohort was previously reported by Vidal et al., 2015 [37], of which we included 13 patients with CD and 7 non-IBD control individuals. Patients who received antibiotic treatment within one month prior to the colonoscopy were excluded.

### Ethics approval

The study was approved by the Institutional Review Board of Clínica Las Condes, Faculty of Medicine, Universidad de Chile; Ethics Committee of the Northern Metropolitan Health Service, Santiago, Chile; and the Balearic Islands’ Ethics Committee, Spain. Study participants provided written informed consent before entering the study. All records and information were kept confidential and all identifiers were removed prior to analysis.

### Nucleic acid extraction, 16S rDNA amplification and pyrosequencing

Total DNA was extracted from biopsy samples using the E.Z.N.A. DNA/RNA Isolation kit (Omega-Bio-Tek) following the manufacturer’s recommendations. The rRNA 16S gene was amplified from the extract using the universal primers GM3 (5′-AGAGTTTGATCMTGGC-3′) and 907r (5′-CCGTCAATTCMTTTGAGTTT-3′). A nested PCR with the amplified product was performed using 454 primers. The primer sequences for the nested PCR are listed in Table S4. This purified product was sequenced using the 454 GS-FLX+ platform (Macrogen, Seoul, South Korea). The datasets of Spanish subjects generated during the current study are available in the ENA sequence repository under the project accession numbers PRJEB6107 and ERP005574.

### Sequence trimming, chimera verification and OTU grouping

Data were processed using a Mothur pipeline [81]. We eliminated low-quality sequences, defined as: sequences under 300 bp, those with a window size and average quality score of 25, a maximum homopolymer of 8 nucleotides, without ambiguities and reading mismatches with barcodes primers. Chimeras were eliminated using Chimera Uchime implemented in Mothur. Sequences were grouped into OTUs with 99% [37] identity using the UCLUST package in QIIME [82]. The most abundant reading of each OTU was chosen as its representative.

### Phylogenetic affiliation and OPU determination

Representative OTUs, grouped into OPUs in a previous study [37] on samples from Spanish patients, were added to the LTP111 database. These samples were aligned using SINA [83, 84] and incorporated into ARB [38] using the maximum parsimony model. The sequences were grouped into OPUs based on a manual inspection of their genealogy.

### Compositional, comparative, and statistical analysis

For the statistical analysis and visual exploration, the microbiome data were uploaded to the MicrobiomeAnalyst server [85]. An alpha diversity analysis was calculated based on observed OPUs (Richness) and Shannon Index (Diversity). Beta diversity was calculated based on the principal coordinate analysis (PCoA) using the Bray-Curtis dissimilarity. A hierarchical cluster analysis based on the relative abundances of 608 OPUs was performed using the Bray-Curtis dissimilarity metric and Ward’s linkage. Heat maps showing OPU abundances were drawn using the gplots package [86] in R [48].

Statistical differences in the abundance of specific taxa between groups were determined using the Kruskal-Wallis test followed by Dunn’s multiple comparisons test. The variation in the read counts and relative abundances across patients can hamper and bias the identification of disease-associated taxa [40] Indeed, as the taxonomic level decreases, there are fewer taxonomic units and therefore, the subject-to-subject variation can lead to the loss of statistical power. To overcome this problem, logarithmic transformation has been implemented in several microbiome studies, allowing better association analyses compared to the linear scale [40, 87]. Consequently, analysis at family level and below were made on data that was transformed using the log(X+1) method.

To determine differentially abundant taxa in different groups, a linear discriminant analysis effect size (LEfSe) [88] was performed in the MicrobiomeAnalyst server [85]. For this, a p-value cut off at 0.05 and linear discriminant analysis (LDA) score threshold 2.0 were used. The LEfSe is an algorithm that detects species and functional characteristics that are differentially abundant between two or more environments with statistical significance, effect relevance, and biological consistency. The LEfSe uses the LDA to estimate the effect size of each differentially abundant feature. To explore the potential interactions among OPUs, a correlation network analysis was performed by using the SparCC algorithm [52].

### Bacterial biomarkers and classifier validation

We defined as biomarkers those OPUs identified by the LEfSe algot which were present in more than half of the samples of the corresponding group (i.e., dysbiosis vs. eubiosis and dysbiotic UC patients vs. dysbiotic CD patients). In order to evaluate the performance of each indicator OPU in discriminating between groups, we first performed a receiver operating characteristic (ROC) curve analysis in the easyROC server [89]. Next, we selected the best discriminating OPUs and implemented several classification algorithms, including Naive Bayes (NB), Random Forest (RF), Logistic Regression (LR) and Support Vector Machine (SVM) and Neural Network (NN) using the Orange data mining suite, V.3.27.0 (http://orange.biolab.si) [90]. NB is a generative model, RF is an ensemble method using decision trees, whereas SVM, LR and NN are discriminative models. We used 70% of samples as the training set, and the rest were used as the test set with the five-fold cross-validation method. The results of the cross-validation (classification accuracy, sensitivity, and specificity) were registered and depicted by ROC curves.

## ACKNOWLEDGMENTS

We thank the National Laboratory for High Performance Computing, NLHPC (ECM-02), for sharing their server facilities, and Mauricio Cerda from the Center of Medical Informatics and Telemedicine (CIMT) at the University of Chile for his technical assistance. We thank Dr. Helen Lowry for her careful review of the manuscript and helpful discussions.

This study was supported by Fondo Nacional De Desarrollo Científico y Tecnológico FONDECYT grant 1161161 to R. Vidal, CONICYT-PCHA/2014-21140975 fellowship to N. Chamorro, FONDECYT 1120577 and 1170648 to Hermoso MA and the Spanish Ministry of Economy projects CLG2015 66686-C3-1-P to Rosselló-Ḿora R., as well as funds from the the European Regional Development Fund (FEDER) and NSF Dimensions in Biodiversity grant OCE-1342694. Support was also provided by a Millennium Science Initiative grant from the Ministry of Economy, Development and Tourism to Paredes-Sabja D.

## CONFLICT OF INTEREST

Funding institutions did not influence the study design or interpretation of the result. Moreover, the authors declare that they have no competing interests.

## SUPPLEMENTARY MATERIAL

**Supplementary Figure 1**. Evaluation of OPUs as diagnostic biomarkers in IBD.

**Supplementary Table 1.** Clinical and demographic features of the IBD patients and control individuals enrolled in this study

**Supplementary Table 2.** Read distribution, OTU and OPU reading analysis, characterized by patient group and country

**Supplementary Table 3**. Bacterial clades that differed between IBD patients with dysbiosis and without dysbiosis

**Supplementary Table 4**. Primer sequences used by nested PCR

## Notes

### Competing Interest Statement

The authors have declared no competing interest.

