## Supplementary Figure 1 for "Landscapes and bacterial signatures of mucosa-associated intestinal microbiota in Chilean and Spanish patients with inflammatory bowel disease"

^5^ Programa Enfermedad Inflamatoria Intestinal. Servicio de Gastroenterología, Clínica Las Condes, Santiago, Chile

^6^ Gastroenterología, Clínica Universidad de Los Andes, Santiago, Chile

^7^ Microbiota-Host Interactions and Clostridia Research Group, Departamento de Ciencias Biológicas, Facultad de Ciencias de la Vida, Universidad Andrés Bello, Santiago, Chile.

^8^ ANID - Millennium Science Initiative Program - Millennium Nucleus in the Biology of Intestinal Microbiota, Santiago, Chile.

^9^ Department of Biology, Texas A&M University, College Station, TX, 77843, USA

^10^ Department of Gastroenterology and Palma Health Research Institute, Hospital Universitario Son Espases, Palma de Mallorca, Spain.

^11^ Grupo de Microbiología Marina, IMEDEA (CSIC-UIB), Esporles, Illes Balears, Spain

^12^ Instituto Milenio de Inmunología e Inmunoterapia, Facultad de Medicina, Universidad de Chile, Chile

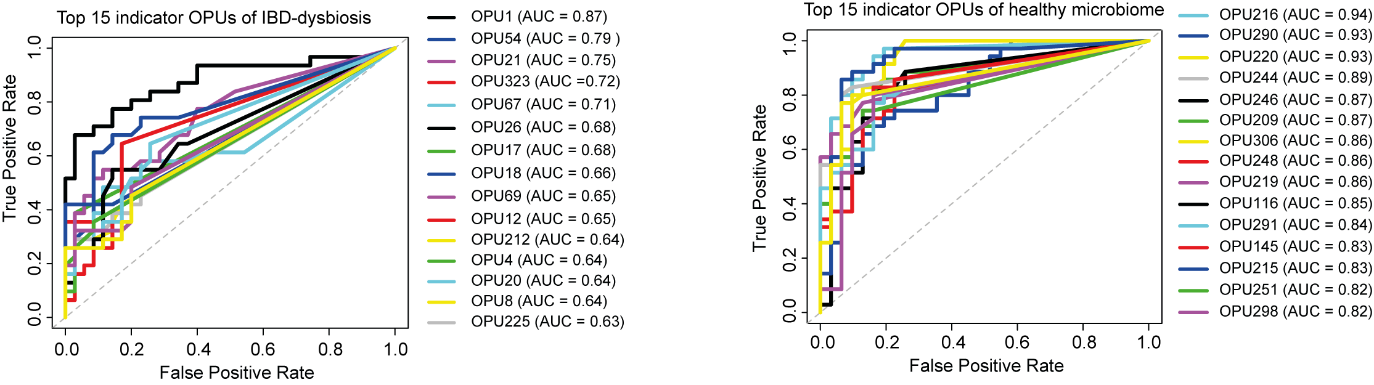

**FIGURE SUPPLEMENTARY 1. Evaluation of OPUs as diagnostic biomarkers in IBD.** OPUs identified by the LEfSe algorithm as differentially abundant in dysbiotic IBD patients or controls were also evaluated as biomarkers to discriminate between dysbiosis and eubiosis. The receiver operating characteristics (ROC) analysis was performed using the easyROC server [1]. AUC, Area under the curve.
